# More sustainable vegetable oil: balancing productivity with carbon storage opportunities

**DOI:** 10.1101/2021.05.14.444195

**Authors:** Thomas D Alcock, David E Salt, Paul Wilson, Stephen J Ramsden

## Abstract

Intensive cultivation and post-harvest vegetable oil production stages are major sources of greenhouse gas (GHG) emissions. Variation between production systems and reporting disparity have resulted in discordance in previous emissions estimates. To assess systems-wide GHG implications of meeting increasing edible oil demand, we performed a unified re-analysis of life cycle input data from diverse oil palm, soybean, rapeseed, and sunflower production systems, from a saturating search of published literature. The resulting dataset reflects almost 6,000 producers in 38 countries, and is representative of over 74% of global vegetable oil production. Determination of the carbon cost of agricultural land occupation revealed that carbon storage potential drives variation in production GHG emissions, and indicates that expansion of production in low carbon storage potential land, whilst reforesting areas of high carbon storage potential, could reduce net GHG emissions whilst boosting productivity. Nevertheless, there remains considerable scope to improve sustainability within current production systems.

## Introduction

From around 800,000 years ago, up to the year 1800, atmospheric carbon dioxide (CO_2_) concentrations averaged around 225 parts per million (ppm)^1^. Despite regular fluctuations, coinciding with ice ages and interglacial periods, concentrations never rose above 300 ppm during this time. However, since the early 1900s, atmospheric CO_2_ concentrations have failed to drop below 300 ppm^2^. In every year since 2015, they have remained above 400 ppm, 70-80% higher than pre-industrial concentrations^3,4^. The Intergovernmental Panel on Climate Change (IPCC) stated that the dominant cause of global warming since 1950 has been anthropogenic contributions to greenhouse gas (GHG) emissions^5^. As the human population has grown, food production has risen markedly; today, food supply chains are responsible for 26% of all GHG emissions^6^. As we strive to provide greater amounts of nutritious food to over 800 million currently undernourished people^7^, whilst meeting additional demand as the population continues to grow^8^, carefully targeted global food systems interventions are required to limit the effects of increased food production on planetary health.

Vegetable oils are a major source of dietary polyunsaturated fatty acids^9^, and are a crucial component of wide-ranging cuisine. Steadily increasing demand for edible oil over at least the last 60 years has led to increased oil crop production through expansion of cultivation area^10,11^ (**Figure 1**) and intensifying production practices^12^. Since 2014, oil crops have inhabited over 300 million hectares (ha) globally, approximately 19% of total cropped land (excluding pasture)^10^. Strikingly, over 85% of the world’s vegetable oil is produced by just four crops: oil palm, soybean, rapeseed and sunflower^10^, which are distributed across a range of climate zones. Clearing of native vegetation to meet growing demand for these crops^13^ represents a considerable source of GHG emissions, further exacerbated by intensive cultivation and post-harvest processing^14^. Estimates of the associated GHG emissions using life cycle assessment (LCA)^15^ have been widely reported. Significant variation exists between assessment results, some of which reflects regional variation and varied production practices^6^. However, despite genuine differences between systems, it is likely that considerable variation between studies is a result of non-harmonious reporting. Additionally, functional units used in analyses vary between studies, further complicating meaningful comparison.

**Figure 1.**
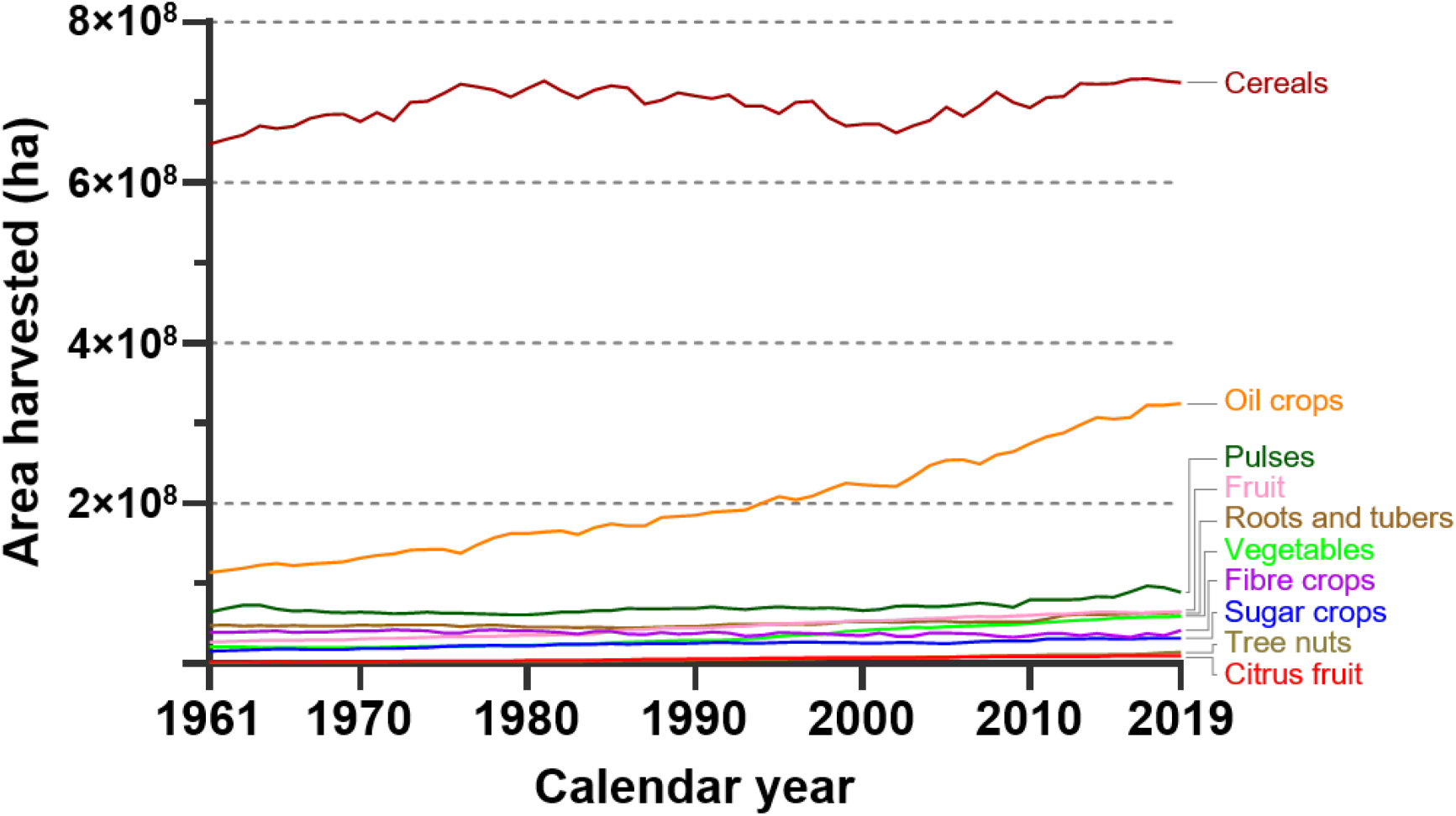
Global area harvested of ten major crop groups in hectares (ha), from 1961 to 2019 (inclusive). Data from FAOSTAT Statistical Database^10^.

Here, we present the results of a harmonised re-analysis of GHG emissions from palm, soybean, rapeseed, and sunflower oil production, using raw input and emissions source data obtained through a saturating search of published literature. The resulting dataset represents more than 74% of global vegetable oil production systems. Such an approach has previously proven useful for deducing variation in life cycle GHG emissions between diverse food production systems^6^. We combine this with a systems-wide analysis of the carbon costs of agricultural land occupation, following carbon storage opportunity principles^16^: these principles allow both recent land use changes and the choice to continuously occupy ancestrally cleared land to be considered equally. This unified, systematic analysis reveals the carbon impacts of vegetable oil production decisions at a global scale, and provides information on how to reduce GHG emissions, both within and between crop systems.

## Results

### Building the global oil crop emissions database

We modelled life cycle GHG emissions resulting from vegetable oil production by combining land use emissions analyses with a harmonised re-analysis of raw data obtained from a saturating search of published literature. We specifically focussed on vegetable oil derived from oil palm, soybean, rapeseed and sunflower, which account for over 85% of global vegetable oil production. We performed initial literature searches on 13^th^ February 2020 using eight bibliographic databases (**Supplementary Data 1**), and additionally monitored Web of Science email alerts, based on initial search terms, throughout 2020. A total of 2,814 unique literature sources were identified for potential inclusion, of which 253, published between the years 2000 and 2020, were retained for quantitative analysis after assessment against nine inclusion criteria (**Supplementary Data 2-7**). The resulting literature set reflects almost 6,000 producers in 38 countries, and is representative of 74.1% of global vegetable oil production (**Supplementary Data 8**). The literature set contains sources corresponding to major production regions for oil palm (South-East Asia), soybean (China, USA, Brazil, Argentina), rapeseed (Canada, Germany, India), and sunflower (Ukraine)^10^. However, no relevant literature records were identified for rapeseed production in China, or sunflower production in Russia, despite these being the second largest producers of rapeseed and sunflower oil, respectively. It is possible that sources exist for these production systems in non-English languages, which were not included in this analysis.

We assessed oil production GHG emissions from crop cultivation through to oil refining (**Figure 2**). We manually extracted production system meta-data, output, material and energy input and direct emissions source data from the literature set, and used these to populate custom life cycle databases. Extraction of raw input data, rather than reported emissions values, was crucial for achieving consistent and comparable results, due to variation in emissions analysis methodology between studies. Where multiple production systems, distinct in crop type, geography and/or production type, were included within single literature sources, we treated each as individual records. Data from a total of 439 records across crops were compiled (**Supplementary Data 8**). We aggregated input data into 22 cultivation and 27 post-harvest stage data-items (**Supplementary Data 9**), to facilitate meaningful comparison between records. We collected data for an additional 33 data items for palm oil, to reflect site preparation, seedling production, and non-productive stage inputs only relevant to oil palm. We then consolidated records into specific production systems, based on geography and cultivation/processing methods, to as far as possible eliminate error and reporting gaps present in individual records. Ninety-six distinct vegetable oil production systems are thus represented here. Finally, we used systems life cycle input data (**Supplementary Data 10-13**) to calculate associated GHG emissions, using data-item specific emission factors from a database compiled as part of this study (**Supplementary Data 14**). Emissions are reported here as kg carbon dioxide equivalent (CO_2_*e*) per kg refined oil, allocated to vegetable oil in each system as a proportion reflective of the economic value of the oil fraction of crop output. Non-allocated emissions data are additionally presented (**Supplementary Data 15-18)**.

**Figure 2.**
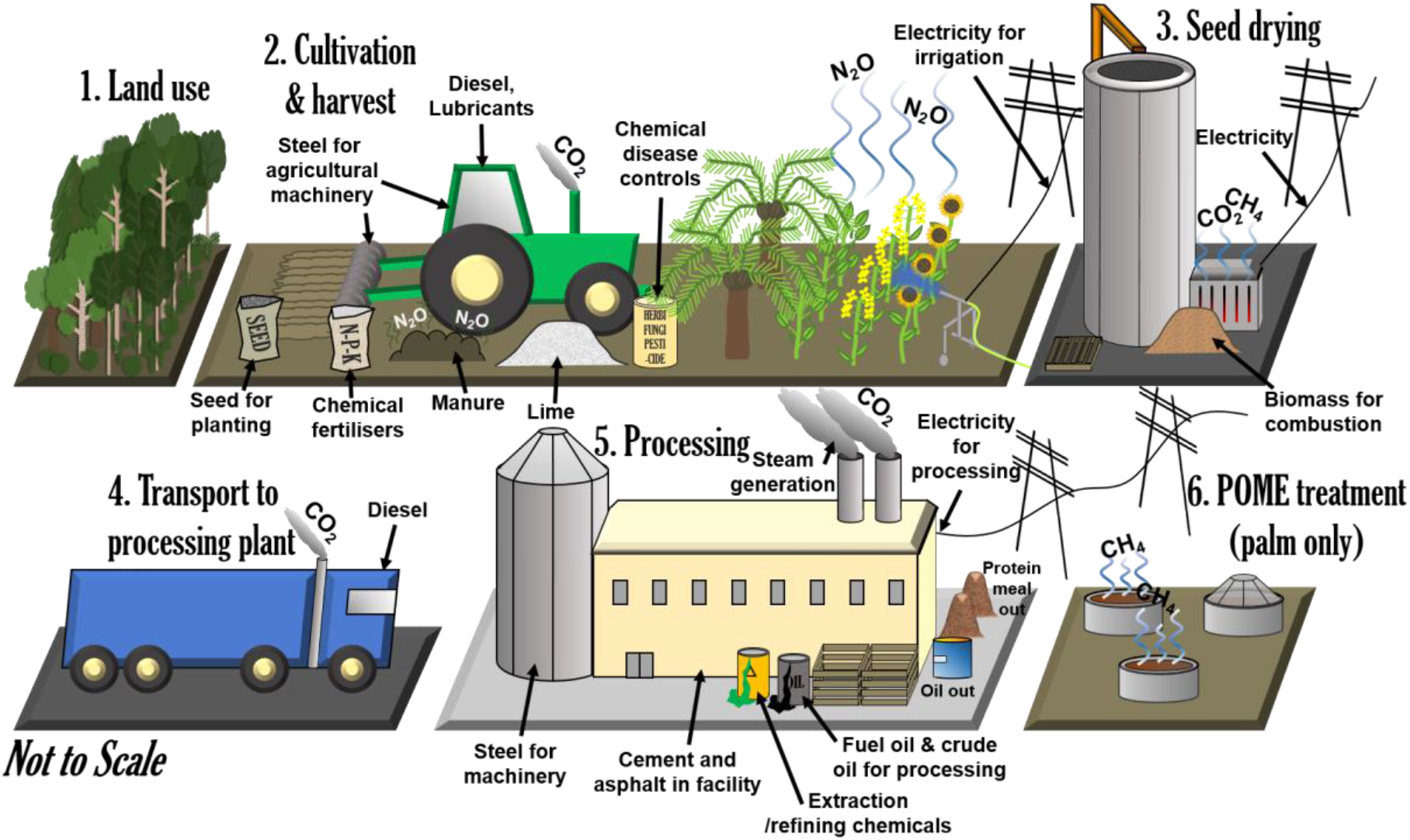
System boundaries of the harmonised reanalysis of life cycle greenhouse gas emissions from vegetable oil production. Major inputs and emission sources indicated.

### Carbon storage potential as a land use indicator

Emissions resulting from land use are often omitted from life cycle assessments, or alternatively, only recent land use changes are generally considered^17^. Failure to assign land use costs to crops grown on ancestrally cleared land could result in intergenerational inequity. For instance, most land clearance for agriculture in Europe took place prior to the 1800s, whereas cropland in various developing regions, including Latin America and SE Asia, has been expanding steadily over the last 100 years^18^. Thus, if only recent land use changes are considered, crops grown in what have become higher income countries may be assigned lower life cycle GHG emissions than those grown in developing countries. Whilst only minimal carbon stock changes might be expected from continuous agricultural occupation of ancestrally cleared land, it is likely that such land could store more carbon if it were set aside for regeneration of native vegetation.

To overcome land use change metric inequity, we modelled the impacts of agricultural land occupation here using carbon storage opportunity principles, as described in detail by Searchinger *et al.*^16^. In essence, we explicitly acknowledge that for each year of continuous agricultural land use, an opportunity to sequester carbon from the atmosphere is lost. We implemented this by comparing the carbon stock of native vegetation and soil in a given area with the carbon stock of vegetation and soil in the same area used for crop production^19^. The difference in carbon stored between the two systems can be considered a carbon storage opportunity cost, if the land use with the lower carbon storage potential is maintained. Considering land use in these terms enables more accurate comparison of the carbon costs of agricultural land occupation, irrespective of if, or when, land use change occurred.

### High yielding crops for lower land use impacts

The environmental impacts of land use can be balanced by productivity. If a given system can produce large amounts of food per unit area, it may be more efficient to use that land for agriculture, freeing up space elsewhere to store carbon more effectively. We consider two vegetable oil production systems from our analysis in **Figure 3**. The presented systems are representative of approximately 40% of global palm oil, and 12% of global rapeseed oil production, respectively. Native tropical rainforest in SE Asia has a total carbon stock of 290 tonnes per hectare, whereas one hectare of oil palm has a carbon stock of 136.6 tonnes (**Figure 3a**). Deforesting one hectare of rainforest to grow oil palm would therefore represent a carbon storage opportunity cost of 153.4 tonnes, whilst yielding 3,657 kg refined oil per year. Forest in Germany has a carbon stock of 179 tonnes per hectare and one hectare of rapeseed 99.4 tonnes (**Figure 3b)**. Whilst the carbon storage opportunity cost between these land uses is only 79.6 tonnes, rapeseed is less productive than oil palm: 2.65 hectares are required to provide the same quantity of oil per year as one hectare of oil palm. The carbon storage opportunity cost between 2.65 hectares of temperate forest and rapeseed is 210.7 tonnes, higher than that of the oil palm system (153.4 tonnes). We alternatively compare the total carbon stocks of these two scenarios assuming that one offsets the other (**Figure 3c**). In Scenario 1, we dedicate 2.65 hectares to rapeseed production in Germany, sparing one hectare of land in SE Asia. Total carbon stored among all vegetation and soils in this scenario is 553 tonnes. In Scenario 2, we dedicate one hectare to oil palm production in SE Asia, sparing 2.65 hectares of temperate forest in Germany. The carbon stored in the latter scenario is higher (610 tonnes), suggesting that this is the more efficient use of land for oil production. However, it is stressed that for oil palm production to result in more carbon stored overall, the corresponding area used for rapeseed production must actively be dedicated to regeneration of forest. Rapeseed is also a larger source of animal feed than oil palm, which could offset animal feed production elsewhere, potentially shifting the balance of results presented. Note that this metric only considers GHG emissions, and not the impact of land use on other sustainability indicators such as biodiversity.

**Figure 3.**
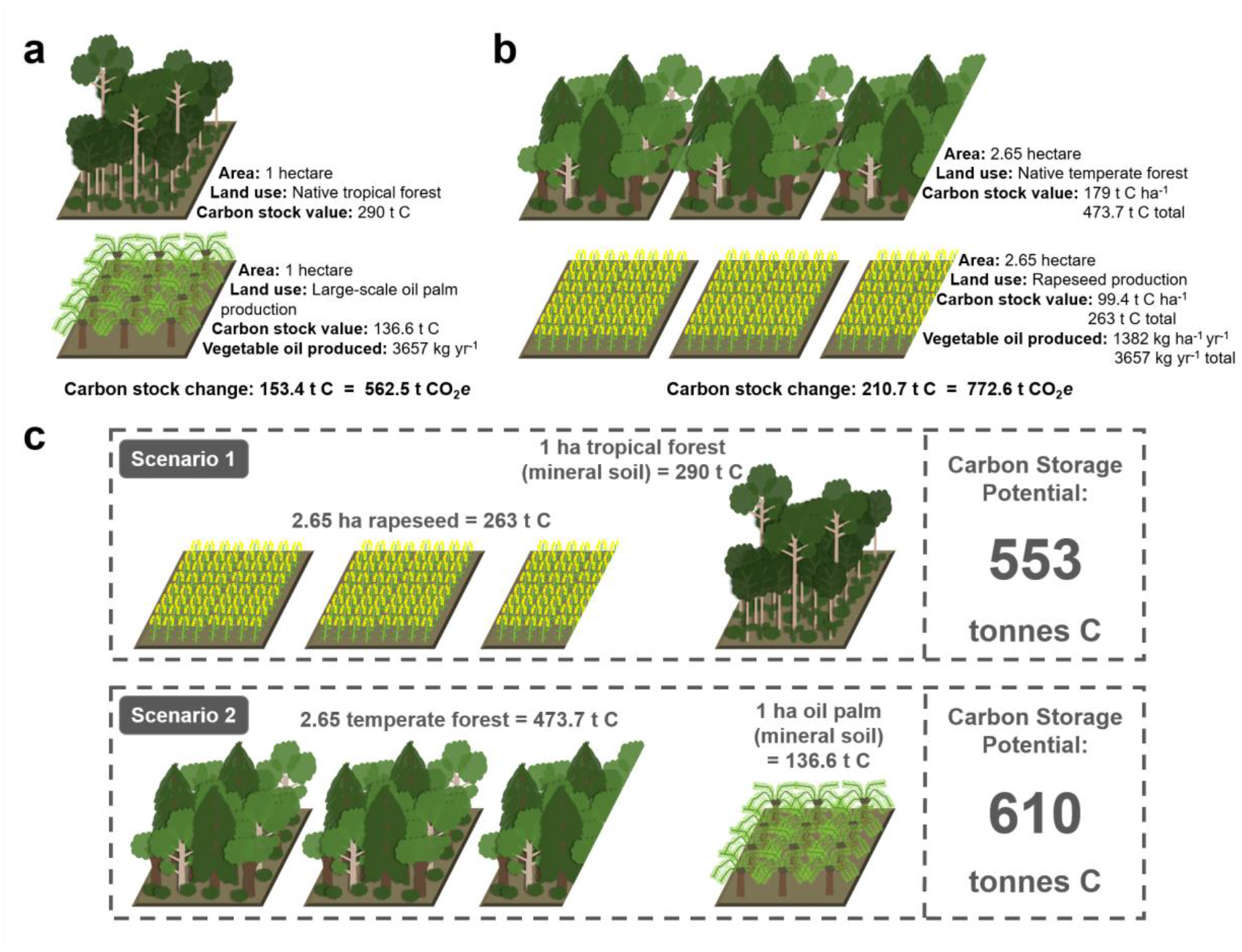
Effect of land use on carbon storage. **a**: Carbon stored in one hectare (ha) of native tropical forest and a large-scale palm oil plantation in South-East (SE) Asia on mineral soils. **b**: Carbon stored in temperate forest and a conventional rapeseed field in Germany in an area which yields the same quantity of vegetable oil as one hectare of oil palm. **c**: Total carbon storage potential of two vegetable oil production scenarios that result in the same quantity of vegetable oil. In Scenario 1, rapeseed cultivation is favoured, allowing tropical forest in SE Asia to be maintained or reforested. In Scenario 2, oil palm cultivation is favoured, allowing temperate forest in Germany to be maintained or reforested. Production systems shown here were selected as representative examples of each crop, based on life cycle emissions from each falling close to the crop specific median, and on a relatively large number of literature records available for each.

### Low native carbon stock land for sustainable oil production

We applied carbon storage opportunity losses/gains between native and agricultural land uses as a carbon penalty/credit to each vegetable oil production system. This was expressed as CO_2_*e*, amortised permanently over 100 years (**Supplementary Data 19-22**). Only one vegetable oil production system in this study was associated with a carbon storage opportunity gain: areas of Canada for which the land cover is native temperate steppe store 11.75 tonnes less carbon per hectare than the same land used for no-till rapeseed production. This is a result of low initial carbon stocks in native biomass, combined with high agricultural inputs including manure addition to the soil, which can build soil carbon stocks^19^. Allocated to refined oil, this carbon storage opportunity gain corresponds to a 0.46 kg CO_2_*e* reduction in life cycle emissions per kg rapeseed oil produced in this system. Similarly, soybean and rapeseed grown in the USA, and soybean, rapeseed and sunflower grown in Iran, can have low associated GHG emissions, resulting from low native vegetation and soil carbon stocks. Land use emissions from other rapeseed production systems ranged from 0.90 to 4.91 kg CO_2_*e* per kg refined oil (**Supplementary Data 21**), whilst sunflower land use emissions fell within a similar range from 0.99 to 6.90 kg CO_2_*e* per kg refined oil (**Supplementary Data 22**). Land use emissions for most soybean systems ranged from 0.36 to 5.53 kg CO_2_*e* per kg refined oil, but two systems, corresponding to production in South Africa and Nigeria, had higher emissions of 7.05 and 15.32 kg CO_2_*e* per kg refined oil, respectively (**Supplementary Data 20**). Meanwhile, land use emissions from palm systems fell into two groups, with emissions from oil palm grown on mineral soils ranging from 0.94 to 1.67 kg CO_2_*e* per kg refined oil, and on peat soils from 24.41 to 28.69 kg CO_2_*e* per kg refined oil (**Supplementary Data 19**). Unsurprisingly, yield was negatively correlated with soybean (df = 25; *R*^2^ = 0.51; *P* < 0.001), rapeseed (df = 35; *R*^2^ = 0.14; *P* = 0.023) and sunflower (df = 21; *R*^2^ = 0.60; *P* < 0.001) land use emissions: greater productivity per hectare could effectively spare land elsewhere for regeneration of native land cover. Soybean (df = 25; *R*^2^ = 0.42; *P* < 0.001), rapeseed (df = 35; *R*^2^ = 0.12; *P* = 0.038) and sunflower (df = 21; *R*^2^ = 0.33; *P* = 0.006) land use emissions were also positively correlated with native vegetation carbon stocks, whereas oil palm land use emissions were very much a product of native soil type (df = 11; *R*^2^ = 0.99; *P* < 0.001; all simple linear regressions).

### Current production systems not optimised for sustainability

We combined systems’ land use emissions data with life cycle GHG emissions assessed through comprehensive re-analysis of published data. Variation in total vegetable oil production emissions across global production systems is presented in **Figure 4**. For each crop, production emissions are fitted against the contribution of each system to global production. Based on the economically allocated dataset, total GHG emissions resulting from vegetable oil production in the across-crop median production system are 3.76 kg CO_2_*e* per kg refined oil. The across-crop median system also forms the median palm oil production system, which is unsurprising since oil palm is currently the largest source of vegetable oil globally^10^. Thus, median system emissions from palm oil production are the same as the across-crop median. Median soybean oil GHG emissions are higher than the global median: 4.25 kg CO_2_*e* per kg refined oil. Median rapeseed and sunflower oil GHG emissions are lower than the global median: 2.49 and 2.94 kg CO_2_*e* per kg refined oil, respectively. Life cycle GHG emissions from palm oil production are dependent on soil type and choice of methane capture technology. Palm oil produced on peat soils is associated with the greatest life cycle GHG emissions across all crops. In contrast, capturing methane emitted by palm oil mill effluent (POME) can reduce emissions by over 50% within certain production systems, which could take palm oil life cycle GHG emissions below rapeseed and sunflower median emissions values. However, despite the clear benefit of methane capture technology, it is currently only adopted by approximately 5% of oil palm mills^20^. For soybean and rapeseed, the lowest emissions are associated with vegetable oil production systems on land with low native carbon stocks, specifically in the USA and in no-till systems in Canada. Emissions resulting from rapeseed production in conventional tillage systems in Canada are more than twice as high as in no-till systems, as a result of differences in carbon stored in soils between each system (**Figure 4**; **Supplementary Data 19-22**).

**Figure 4.**
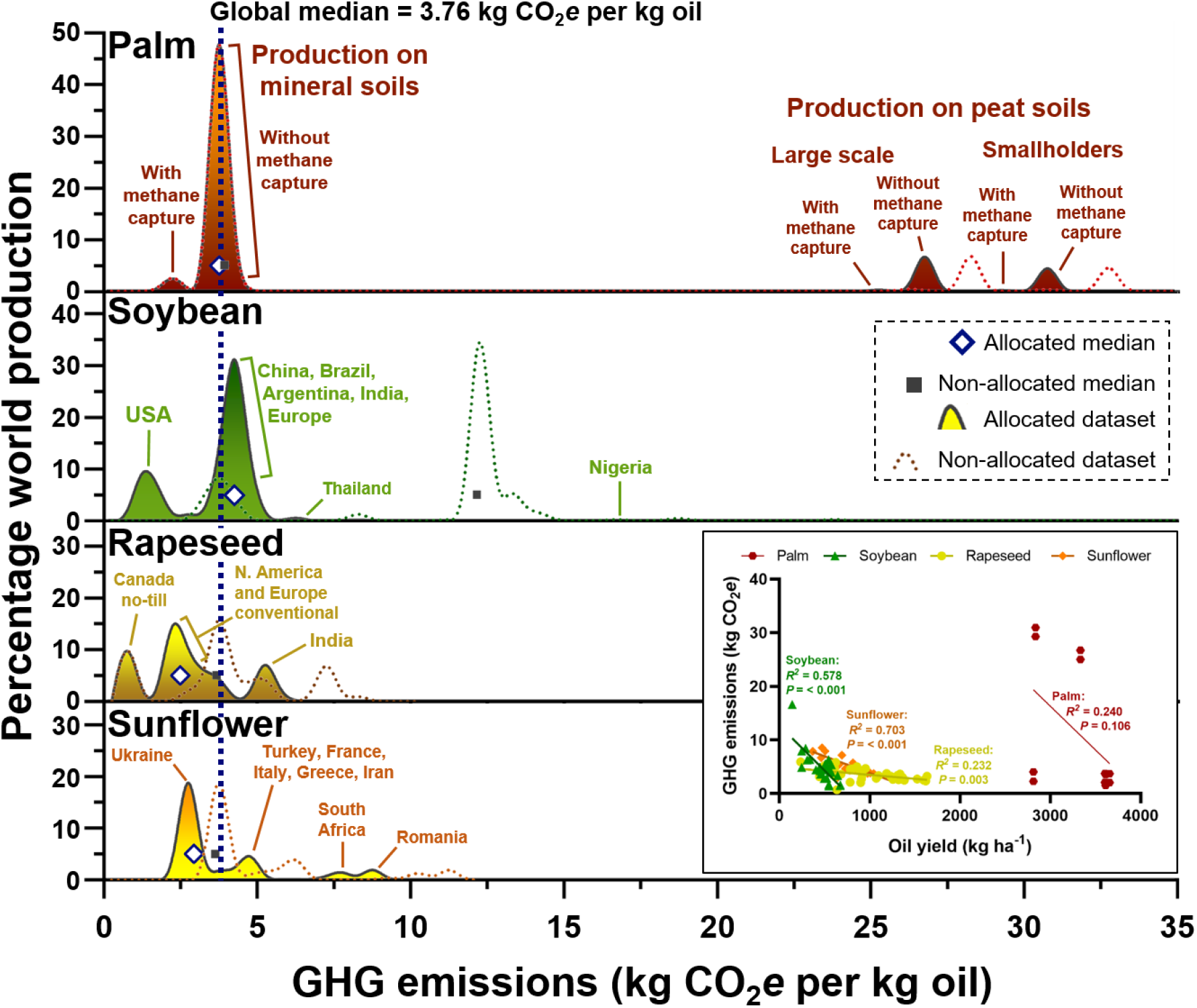
Greenhouse gas (GHG) emissions as CO_2_ equivalent (CO_2_*e*) resulting from global palm (12 systems; 147 records), soybean (26 systems; 106 records), rapeseed (36 systems; 128 records) and sunflower (22 systems, 58 records) oil production systems. Emissions allocated by economic value to oil shown as filled curves, with non-allocated emissions shown as dotted curves for reference. The height of each curve represents the percentage of global production from each crop that results in the specified GHG emissions. Median GHG emissions from each crop indicated by white diamonds. Median GHG emissions from all crops combined, weighted by contribution to world vegetable oil production, shown as dashed blue line. Note one data point from non-allocated dataset outside of displayed range for soybean (conventional production in Nigeria; 49.56 kg CO_2_*e* per kg oil). Figure annotated with selection of pronounced production systems for reference, referring to the allocated emissions dataset in each case. Figure inset (bottom right) shows simple linear regressions between oil yield and life cycle GHG emissions for palm (df = 11), soybean (df = 25), rapeseed (df = 35) and sunflower (df = 21) oil production, based on allocated datasets.

The world’s largest producer of sunflower oil, Ukraine, has the production system associated with the second lowest crop-specific GHG emissions. However, within all other crops, it is clear that there is significant scope to reduce GHG emissions (**Figure 4**). This could be achieved through more widespread adoption of emissions-reducing technologies, or through shifting the geographic production range. Soybean, rapeseed and sunflower life cycle GHG emissions are also strongly negatively correlated with yield (**Figure 4 inset**). It follows that if GHG emissions per hectare can stay broadly the same whilst increasing productivity, the total emissions per unit of final product are effectively reduced. A major focus should therefore be on sustainably increasing production on land already occupied by agriculture. However, care should be taken to avoid increasing production through means that result in large amounts of additional emissions. For instance, one might seek to increase yield through applying greater quantities of synthetic nitrogen. However, synthetic nitrogen is associated with almost 6 kg CO_2_*e* per kg applied^21^. Therefore, a better approach might be to identify genotypes with a high yield potential under relatively low nitrogen supply^22^.

### Mitigating emissions through management choices

Whole life cycle emissions range from 0.73 kg CO_2_*e* per kg refined oil for no-till rapeseed oil production in Canada, to 30.96 kg CO_2_*e* for smallholder palm oil production in SE Asia, on peat soils without methane capture technology (**Figure 5**; **Supplementary Data 19-22**). Whilst much of this variation is driven by land use, considerable variation in emissions from other production stages also exists. For instance, soybean cultivation emissions range from 0.27 to 3.89 kg CO_2_*e* per kg oil. The range in cultivation emissions is lower for other oilseeds, but still varies 3.55-fold and 5.75-fold between rapeseed and sunflower production systems, respectively (**Figure 5**). Solutions to reduce production stage emissions are specific for each system. Production of soybean, rapeseed and sunflower in Iran is associated with high emissions from electricity generation, used to power irrigation systems. Similarly, high GHG emissions from seed drying and storage in some regions are a result of high electricity production footprints (**Supplementary Data 14-18**). Reducing electricity requirements for irrigation or seed drying is perhaps unrealistic, but shifting to more sustainable sources of electricity could bring cultivation emissions down^23^. Transport emissions could be reduced by decreasing the distance between cultivation and processing centres, or increasing the fuel efficiency of transport vehicles. The single biggest change that could be implemented to reduce emissions from most palm oil production systems is adoption of POME methane capture technologies. POME is a bigger source of GHG emissions than land use in the median palm oil production system, responsible for 46.5% of life cycle emissions (**Figure 6a**). For all other crops, land use is the dominating source of life cycle GHG emissions (**Figure 6b,c,d**). However, synthetic nitrogen application represents a further major source of emissions, particularly for rapeseed (**Figure 6c**) and sunflower (**Figure 6d**) oil production systems, whilst agricultural diesel use forms the biggest source of non-land-use GHG emissions from both soybean (**Figure 6b**) and sunflower oil production systems.

**Figure 5.**
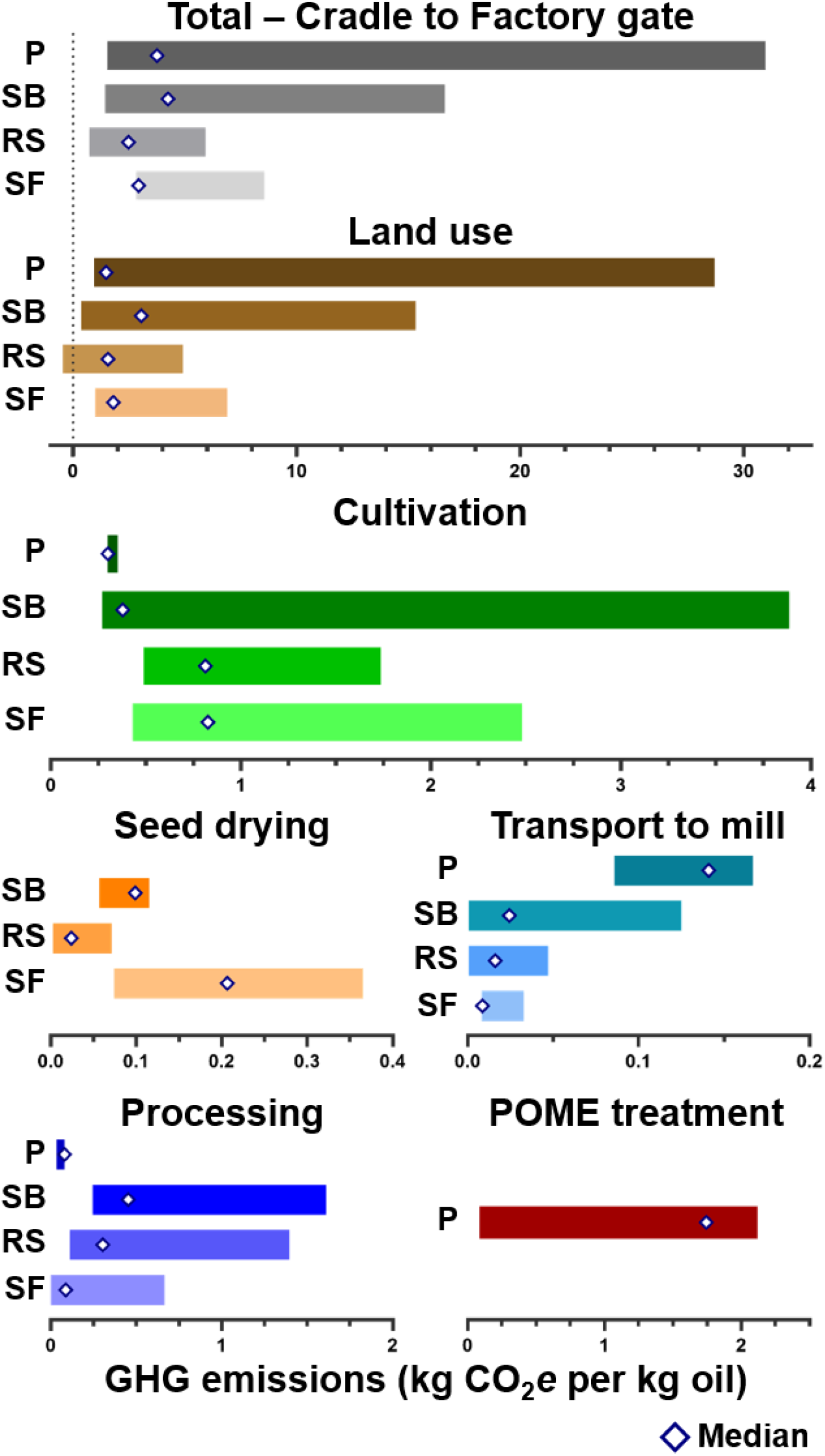
Range of greenhouse gas (GHG) emissions as CO_2_ equivalent (CO_2_*e*) observed across all palm (P; 12 systems), soybean (SB; 26 systems), rapeseed (RS; 36 systems) and sunflower (SF; 22 systems) oil production systems included in this study. Each bar shows full, non-weighted dataset from all systems included in this study, grouped by production stage. Emissions allocated to oil portion of crop system outputs by economic value. Median system GHG emissions per crop per growth stage indicated by white diamonds.

**Figure 6.**
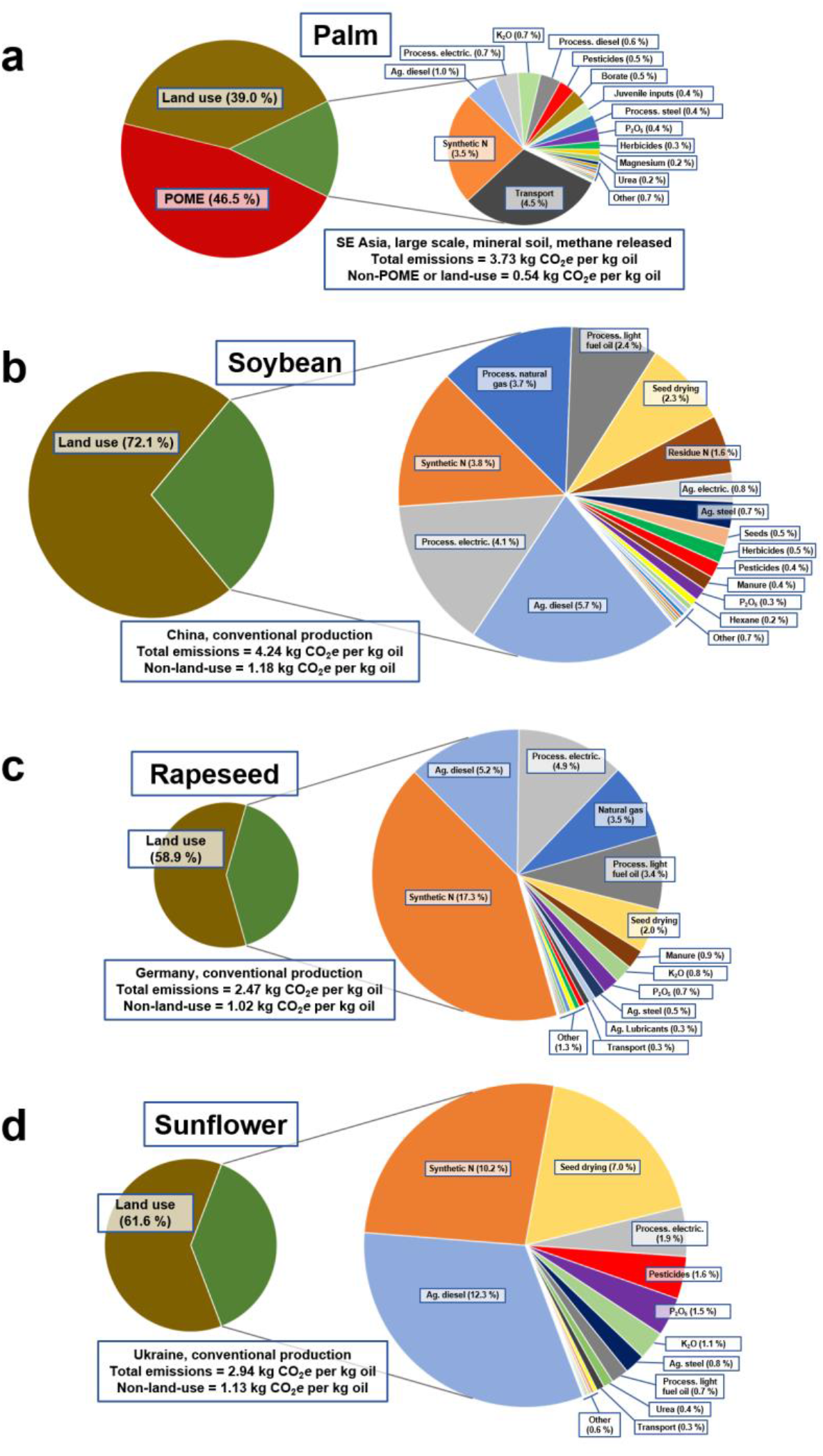
Contribution of input parameters to total life cycle greenhouse gas emissions for specific palm, soybean, rapeseed and sunflower oil production systems. Systems shown are palm oil production in South-East (SE) Asia on mineral soil without methane capture technology (**a**), conventional soybean production in China (**b**), conventional rapeseed production in Germany (**c**), and conventional sunflower production in Ukraine (**d**). Production systems shown here were selected as representative examples of each crop, based on life cycle emissions from each falling close to the crop specific median, and on a relatively large number of literature records available for each. Pie charts on left in each panel scaled to allow comparison between crops, with emissions not caused by palm oil mill effluent (POME) or land use shown in expanded pie charts to the right, again scaled to enable between-crop comparisons.

## Discussion

For at least most of the current decade, vegetable oil demand and production is projected to continue to grow^13,24^. Increasing demand is particularly expected in developing countries in line with rising per capita income, but high consumption in developed regions is also expected to be maintained. Oil palm plantation land area still appears to be growing in SE Asia, and the world production of soybean, rapeseed and sunflower is forecast to continue to grow at a rate of around 1.5% per year^13^. Whilst the use of vegetable oil for biofuel production has fallen out of favour due to sustainability concerns^16,25,26,27^, the demand for used cooking oil as a fuel source is expected to rise significantly^28^. However, despite this being seen as a highly sustainable source of transport fuel compared to first generation biofuel or fossil alternatives^29,30,31^, recent reports suggest that it could in fact be linked to deforestation and thus an additional source of currently under-considered GHG emissions^28,32^. This is partially a result of large imports of used cooking oil from regions where this is often not considered a waste-product but rather a source of animal feed^33^. This requires animal feed to be sourced from elsewhere, leading to increased demand for additional production capacity. Increasing demand for used cooking oil could also incentivise fraudulent sale of fresh oil in its place, reducing the global supply of vegetable oil for other uses^28^. This highlights the importance of producing vegetable oil as sustainably as possible, whilst ensuring that any production or use decisions do not result in increased emissions in other parts of the global food system.

Clearing of native land cover for agricultural expansion can represent a large source of carbon emissions, and should generally be avoided. However, we have shown here that expansion of vegetable oil production in areas of low native carbon stocks or high productivity could, in principle, lead to greater net carbon storage, as long as currently occupied areas with lower productivity, or higher carbon storage potential, are in parallel set aside for regeneration of native land cover. In practice, this would likely require concerted efforts of multiple governments and stakeholders, and perhaps even a global carbon credit system, whereby producers pay for regeneration and maintenance of forests elsewhere. It is also essential to assess alternative sustainability indicators between land uses. For instance, clearing of forest in SE Asia for the production of oil palm represents not only a question of carbon emissions, but also one of biodiversity. Oil palm expansion has been linked to extensive reduction in species richness and abundance across taxa including of insects, birds, small mammals and primates^34^. This must be properly considered before making any global land use change decisions, as it is unlikely for biodiversity to be completely restored to pre-clearance levels in reforested land once lost^35^. It is difficult to see a global sustainability accounting system implemented over the next few years. However, without globally integrated solutions to rising carbon emissions that acknowledge both production system and land use impacts, we are unlikely to reach net zero emissions targets.

## Methods

### Study aim and strategy

The aim of this study was to characterise global systems-wide variation in life cycle greenhouse gas (GHG) emissions resulting from the production of oil palm (*Elaeis guineensis*), soybean (*Glycine max*), rapeseed (*Brassica* spp.) and sunflower (*Helianthus annuus*) derived vegetable oil. This was achieved through a harmonised re-analysis of primary data sources, combined with systems-wide calculations of the carbon cost of agricultural land occupation, as modelled through the concept of carbon storage opportunities^16^. Due to variation in emissions calculations, system boundaries, and functional and time units between studies, extraction of raw emissions source data from the literature, rather than reported emissions values, was crucial for achieving consistent and comparable results. Life cycle input data were used to calculate associated life cycle GHG emissions, reported as CO_2_*e*, based on a custom database of emission factors curated as part of this study (**Supplementary Data 14**). Land use carbon storage indicators were generated taking account of native and crop-specific vegetation and soil carbon stocks. Associated CO_2_*e* values were then additionally assigned to each production system as a function of the difference between native and agricultural land use carbon stocks, permanently amortised over 100 years.

### System boundaries

Each system studied was split into distinct production stages: 1. Land use, 2. Cultivation and harvest, 3. Seed drying and storage (all crops except oil palm), 4. Transport to processing facilities, 5. Processing and refining, and 6. Treatment of palm oil mill effluent (POME; oil palm specific; **Figure 2**). Stages post-refining such as packaging, distribution and use are omitted, due to their limited reporting and highly variable nature. A full list of data items collected within each production stage can be found in **Supplementary Data 9**. The life cycle of oil palm production is considerably different from that of the other crop types included here. Whilst soybean, rapeseed and sunflower are annual crops, sown and harvested within the same twelve months, a single oil palm plantation is generally maintained for around 25-30 years, and includes seedling production and juvenile stages during which time no vegetable oil is output. To account for this, the entire oil palm life cycle was modelled from seedling production to end of productive lifespan per hectare. Resulting GHG emissions were then divided by the total plantation lifespan in years to obtain normalised annual GHG emissions per hectare. Inputs of services such as cleaning, marketing, accounting, and overheads including office space electricity and upkeep were omitted, due to a lack of reporting in studies included in the meta-analysis.

### Functional units

For systems’ modelling and spreadsheet management, energy and material inputs are referred to on a per hectare (ha) basis, since this unit is most relevant to decision making at the cultivation stage. For the purpose of final results’ reporting, the functional unit is defined as one kg of refined vegetable oil, which enables clear comparison of results between crop systems.

### Information sources, search strings and record compilation

To thoroughly extract all relevant literature, eight individual bibliographic databases were consulted. These were Web of Science (all databases), Scopus, PubMed, PubMed Central, Wiley Online Library, SpringerLink, JSTOR, and ScienceDirect. These databases were selected based on their multidisciplinary content, search string capacity and overall performance, as analysed by Gusenbauer and Haddaway^36^. Search strings were formulated to identify studies that concerned oil palm, soybean, rapeseed and/or sunflower in the context of oil production and sustainability. Biofuel/biodiesel was also included in the search strings to incorporate studies which may include data relating to earlier production stages (e.g. cultivation of relevant crops). Search strings varied depending on the required syntax of each bibliographic database, but broadly followed the string used for Web of Science as per below:

#### Palm

((“palm” OR “elaeis guineensis”) AND (“life cycle assessment” OR “life cycle analysis” OR “lca” OR “greenhouse gas emissions” OR “greenhouse emissions” OR “carbon footprint” OR “sequestration” OR “nutrient loss”) AND (“oil” OR “biodiesel” OR “biofuel”))

#### Soybean

((“soy” OR “soya” OR “soybean” OR “soyabean” OR “glycine max”) AND (“life cycle assessment” OR “life cycle analysis” OR “lca” OR “greenhouse gas emissions” OR “greenhouse emissions” OR “carbon footprint” OR “sequestration” OR “nutrient loss”) AND (“oil” OR “biodiesel” OR “biofuel”))

#### Rapeseed

((“rapeseed” OR “canola” OR “rape” OR “oilseed rape” OR “brassica”) AND (“life cycle assessment” OR “life cycle analysis” OR “lca” OR “greenhouse gas emissions” OR “greenhouse emissions” OR “carbon footprint” OR “sequestration” OR “nutrient loss”) AND (“oil” OR “biodiesel” OR “biofuel”))

#### Sunflower

[(“sunflower” OR “helianthus”) AND (“life cycle assessment” OR “life cycle analysis” OR “lca” OR “greenhouse gas emissions” OR “greenhouse emissions” OR “carbon footprint” OR “sequestration” OR “nutrient loss”) AND (“oil” OR “biodiesel” OR “biofuel”)]

Full search strings used for all other databases are included in **Supplementary Data 1** along with number of search results returned for each. Additional searches were performed in Web of Science filtered to only include results from *The International Journal of Life Cycle Assessment* with search strings limited to include only crop identifier terms (**Supplementary Data 1**). In general, searches were directed to scan only text in the title, abstract and in any keywords, since searching in full text records led to too many spurious results. Initial searches were performed on 13^th^ February 2020. However, Web of Science email alerts were set up for each of the full search terms listed above. New publications that were indicated by these alerts were screened *ad hoc* throughout the remainder of 2020, and relevant literature items were added to the respective GHG emissions models where necessary. Thus, the literature included in the meta-analysis described here can be considered to represent the entire set of relevant literature present in the consulted databases from the start of 2000 to the end of 2020. Records were managed in EndNote X9 (Clarivate Analytics, Philadelphia, PA, USA).

### Literature eligibility criteria

Studies were assessed for eligibility for inclusion against nine criteria (**Supplementary Data 2**). These were formulated to fulfil the PRISMA statement reporting guidelines, designed to promote transparent and complete reporting of systematic reviews and meta-analyses^37^. Literature was required to be original and complete, published in English between the beginning of the year 2000 and the end of 2020, and to significantly concern production of oil palm, soybean, rapeseed and/or sunflower over other crops, in a commercially viable setting as opposed to experimental or speculative (e.g. on abandoned quarries), in the context of sustainability. Studies were also required to contain life cycle input data relevant to the system boundaries described above, and to frame their input data in terms of one or more of the functional units used here or enable recalculation into such units based on available data.

### Screening

After removal of duplicates, records were exported using custom output styles to Microsoft Excel for screening. Records were initially screened based on publication year, language and type, then by titles and finally abstract, to quickly exclude irrelevant literature. Full text articles were accessed online for the remaining records. Text, tables, figures and supplementary information were consulted to ensure that only relevant literature was retained for analysis. On occasion, unique records corresponding to the same study and/or dataset were identified, for example where a conference paper was submitted prior to a full journal submission. In these cases, only the most complete or recent record was retained. An overview of the number of sources identified, screened, excluded and retained for analysis is reported for each crop in PRISMA-style flow diagrams^37^ in **Supplementary Data 3-6**.

### Data collection process

Data collection for the meta-analysis utilised custom life cycle input databases managed in Microsoft Excel. Each literature record was given a unique source identifier and apportioned to a unique row within the relevant spreadsheet. Where present in each record, summary information including study location, cultivation practices and oil extraction methods was noted. Relevant data were then identified in tables, figures, text and supplementary information, extracted manually, and used to populate the life cycle input database. The reporting of certain data items was simplified in the database to provide a suitable number of values for comparison. For example, chemical disease/pest controls were grouped into herbicide, insecticide, fungicide and unspecified pesticide items, rather than reporting specific chemicals used. Similarly, fertilisers were grouped into major data items including synthetic nitrogen (N), urea N, manure (total weight), phosphate (as P2O5), and potassium oxide (K2O). Life cycle input data were all expressed per hectare in the initial databases. Data that were expressed in alternative units in the literature were converted using other available data. Study-specific input data were used to perform conversions as much as possible. However, values were assumed in cases where such information was not available, including from average values reported in other relevant literature in the life cycle input database, and as a last resort from online databases such as FAOSTAT^10^. Consistent units were utilised for individual data items, including kg for material inputs, and MJ for energy inputs. Where these were reported differently in literature records, values were converted using consistent conversion ratios e.g. 1 kWh = 3.6 MJ, 1 L diesel = 0.832 kg. A full list of data-items collected, conversion factors used, and assumptions made are reported in **Supplementary Data 23**.

### Assessing risk of bias and record consolidation

It was assumed that reporting bias existed within studies, including variation in included data items, and choice of analysing first-hand production data, survey data, regional average data and/or data from unverified assumptions. Bias was also assumed across studies, including underrepresentation of some systems in the literature. To highlight, and where possible address this, the following measures were taken. For each record, it was noted what kind of system was used to acquire input data. Where this was survey or first-hand production data, the number of participants/farms represented was noted. Records were then consolidated into several production systems, based on geographic production range and cultivation/processing methods, as per Poore and Nemecek^6^. For each data item for each system, the mean of all reported values was then calculated and used as the system standardised value. Where data items relevant to the system boundaries of this study were not reported in individual literature records, cells were generally left blank in the input database. The exception was where it was deemed likely that the true value for a specified category was zero if not reported. For example, if a study reported kg of urea N applied to a field but failed to mention synthetic N, it was assumed that no synthetic N was used. In these cases, zero values were added to relevant cells in the input database. Thus, blank cells were left out of subsequent analyses, and imputed zero values were included, on the assumption that they were representative of within-system variation. This approach enabled most data items to be filled for each system, whilst highlighting the extent to which each system was represented in the literature. Where systems were still missing a value for a given data-item, the value from a highly similar production system was used where possible, otherwise the mean average value of data-item values across all systems was generally calculated and used (**Supplementary Data 23**). Where appropriate, this was weighted by system yield. Finally, the number of records present for each system was compared to global, country-specific production data from FAOSTAT^10^, to identify any disparities between production quantity and scientific reporting incidence.

### GHG emission factors database

To enable calculation of GHG emissions from the life cycle input database, a custom emission factors database was compiled. This comprised estimated carbon dioxide (CO_2_), methane (CH_4_) and nitrous oxide (N_2_O) emissions associated with the manufacture, distribution and use of the energy and material inputs under study here. Collection of emission factors relating to the three gasses individually allowed consistent calculation of CO_2_-equivalent (CO_2_*e*) emission factors, which comprehensively represent Global Warming Potential (GWP). For this study, IPCC AR5 GWP100 conversion factors with climate-carbon feedbacks were used^38^. Emission factors were collected from multiple emissions databases including BioGrace^21^, UK Government GHG Conversion Factors for Company Reporting 2019^39^ the EMEP/EEA air pollutant emission inventory guidebook 2019^40^ and the software GREET 2019 (version 1.3, Argonne National Laboratory, IL, USA), or from literature sources **Supplementary Data 14**. Electricity emissions were calculated using country-specific emission factors to reflect regional variation in electricity generation practices. For some data items, only CO_2_*e* emission factor values were available, many of which were calculated using previous GWP conversion estimates. Where recalculation to AR5 values wasn’t possible, these were retained as a best estimate of the emissions associated with the given factor. Of the gasses under study here, only the CH_4_ conversion factor differs between IPCC AR4 and AR5 (with climate-carbon feedbacks). Hence, for data items for which AR4 conversion factors are used here, it is likely that only minimal error in final emissions calculations exists.

### Modelling land use through carbon storage opportunity

Land use was modelled here using the principle of carbon storage opportunity cost^16^. This compares the carbon stock of native vegetation and soil in a given area, with the carbon stock of vegetation and soil, at equilibrium, of the same area used for production of a given crop. The difference in carbon stored between the two systems can be considered a carbon storage opportunity cost, if the land use with the lower carbon storage potential is maintained. This is balanced by productivity, whereby carbon storage opportunity cost is divided by the quantity of food produced. Carbon storage opportunity forms a multi-use indicator, allowing comparison of carbon storage potentials in native vegetation and soils across geographic ranges, between different land uses in a given area, and between different areas of cropland with contrasting food productivity and/or native carbon stocks^16^. Importantly, it allows for comparison of the carbon cost of agricultural land occupation between crop systems, irrespective of if or when land use change actually occurred.

For each production system, ICPP Climate Zone^41^, soil type^42^ and native land cover^43^ data were sourced and used to infer native and agricultural vegetation and soil carbon stocks from IPCC 2006^44^ values via datasets presented in Flynn et al.^19^. Agronomic input levels were grouped by total N application rates, where rates above 100 kg ha^-1^ were considered high, between 50 and 100 kg ha^-1^ medium, and below 50 kg ha^-1^ low input, and used to infer agricultural soil carbon stocks. Resulting data were fed into carbon stock change calculations in the Excel tool provided by Flynn et al.^19^ to determine differences in stored carbon between native and agricultural land uses. Values were divided by 100 to permanently amortise carbon stock changes over 100 years, and attributed to each crop system **Supplementary Data 19-22**. Multi-cropping within one year and fallow periods were not included in land use calculations, due to limited reporting within literature sources and since these were expected to largely offset one another. Land use calculations here assume constant or growing oil crop production; that additional oil can only be produced on land currently used for oil production or by clearing new land; no spatial limitations; and that regeneration of ancestrally cleared land can restore carbon to native levels within 100 years. The latter assumption is likely to be true for most forest systems^45,46^, but not for peatlands for which restoring carbon stocks may take significantly longer^47^.

### Economic allocation and emissions reporting

For each system, life cycle input data were multiplied by the respective emission factor from the emission factors database, and total life cycle and production stage specific emissions were determined. This was on an annual basis per hectare for soybean, rapeseed and sunflower. For oil palm, productive lifetime emissions values were normalised to an annual basis for comparison, by dividing by the plantation lifetime in years, including non-productive years. Annual life cycle emissions values were then combined with amortised carbon storage opportunity losses/gains, and divided by annual, system specific oil yield, for reporting of life cycle GHG emissions per kg refined oil.

Life cycle GHG emissions are reported as a whole for each crop system, and additionally as a proportion reflective of the economic value of the oil portion of total crop produce. Economic allocation of emissions was determined to be the most suitable method for distinguishing between emissions from different products, as this can reasonably be expected to influence land use decisions for a given area of land. Economic values of crop portions were determined primarily using the World Bank Commodities “Pink Sheet” data^48^ and USDA Oilseeds World Market and Trade reports^49^ (**Supplementary Data 24**). Price data from October 2018 to September 2019 were used, rather than more recent data, in order to avoid impacts of COVID-19 on prices. Allocation was performed separately for each system, based on quantified co-product outputs. Production emissions were allocated proportionately between co-products for all emissions sources with the exception of emissions only relevant to refining of vegetable oil after separation from co-products, which were allocated in full to the oil fraction.

### Figure generation

Figures 1 and 5 were generated in GraphPad Prism 9 (GraphPad Software, San Diego, CA, USA). Figure 4 was generated in OriginPro 2021 (OriginLab Corporation, Northampton, MA, USA), where system specific life cycle GHG emissions estimates were binned into 0.5 kg CO_2_*e* intervals and plotted using using B-Spline line functions, weighted by system contribution to world production (Supplementary Data 25). Individual pie charts in Figure 6 were generated in Microsoft Excel 2016, then manually scaled by represented emissions. Simple linear regression analyses were performed in GraphPad Prism.

## Author contributions

TDA contributed to study design, performed initial literature review, curated greenhouse gas emission databases, performed and contributed to interpretation of data analyses, and led writing of the manuscript. DES, PW and SJR contributed to study design, advised data analysis procedures, contributed to the interpretation of analyses and critically reviewed and edited the manuscript. All authors approve of the final version of the manuscript.

## Data availability

Raw data for all figures can be found in the Supplementary Data.

**The authors declare no competing interests.**

